# Isomeric *O*-methyl cannabidiolquinones with dual BACH1/NRF2 activity

**DOI:** 10.1101/2020.06.16.153981

**Authors:** Laura Casares, Juan Diego Unciti, Maria Eugenia Prados, Diego Caprioglio, Maureen Higgins, Giovanni Apendino, Albena T. Dinkova-Kostova, Eduardo Muñoz, Laureano de la Vega

## Abstract

Oxidative stress and inflammation in the brain are two key hallmarks of neurodegenerative diseases (NDs) such as Alzheimer’s, Parkinson’s, Huntington’s and multiple sclerosis. The axis NRF2-BACH1 has anti-inflammatory and anti-oxidant properties that could be exploited pharmacologically to obtain neuroprotective effects. Activation of NRF2 or inhibition of BACH1 are, individually, promising therapeutic approaches for NDs. Compounds with dual activity as NRF2 activators and BACH1 inhibitors, could therefore potentially provide a more robust antioxidant and anti-inflammatory effects, with an overall better neuroprotective outcome. The phytocannabinoid cannabidiol (CBD) inhibits BACH1 but lacks significant NRF2 activating properties. Based on this scaffold, we have developed a novel CBD derivative that is highly effective at both inhibiting BACH1 and activating NRF2. This new CBD derivative provides neuroprotection in cell models of relevance to Huntington’s disease, setting the basis for further developments *in vivo*.

## INTRODUCTION

Cells are continuosuly exposed to reactive oxygen species (ROS) from both exogenous and endogenous sources. To counteract the potential harmful effects of ROS, cells have evolved constitutive (housekeeping) as well as inducible antioxidant systems that regulate redox homeostasis. The master regulator of the inducible antioxidant responses is the transcription factor NF-E2–related factor 2 (NRF2), a member of the cap‘n’collar basic region leucine zipper (CNC-bZip) transcription factor family. NRF2 binds to Antioxidant Response Elements (ARE) in the promoters of its target genes, many of which encode antioxidant and other cytoprotective proteins, and triggers their expression [1]. BTB And CNC Homology 1 (BACH1) is a transcriptional repressor, also belonging to the CNC-bZip family, that competes with NRF2 for binding to ARE sites located at the promoter of oxidative stress responsive genes, such as heme oxygenase 1 (*HMOX1*) [2].

In homeostatic conditions, NRF2 binds to its negative regulator Kelch-like ECH-associated protein 1 (KEAP1) in the cytoplasm, and is targeted for ubiquitination and subsequent proteasomal degradation by the Cullin3-containing E3-ligase complex [3]. When cells are exposed to oxidants or electrophiles, KEAP1 is disabled, allowing for accumulation and translocation of de novo synthesized NRF2 to the nucleus [4] while BACH1 is exported out of the nucleus and degraded [5]. Only a subset of NRF2 target genes are regulated by BACH1, *HMOX1* being the best characterized one. It encodes heme oxygenase 1 (HMOX1), an enzyme which catalyzes the rate-limiting reaction in heme degradation [6]. It is generally accepted that BACH1 and NRF2 work together to regulate the expression of *HMOX1*, with the negative effect of BACH1 dominating over the positive effect of NRF2. This means that BACH1 needs to be displaced from the *HMOX1* promoter in order for NRF2 to bind and induce its expression [7]. However, we and others have shown that *HMOX1* induction can be BACH1-dependent, but NRF2-independent [8] [9], suggesting that the regulation of *HMOX1* is most likely cell type- and context-dependent.

HMOX1 has anti-inflammatory properties [10] [11], and its knockout in mouse models leads to the development of chronic inflammation [12] [13]. The antioxidant role of *HMOX1* is largely attributed to its capacity to degrade heme into biliverdin, which is rapidly converted to the antioxidant bilirubin, free iron, and carbon monoxide [14] [15]. Furthermore, various studies have shown that overexpression of *HMOX1* protects cells from oxidative damage and neurotoxicity [11][16]. Oxidative stress and inflammation in the brain are two key hallmarks of neurodegenerative diseases (NDs) such as Alzheimer’s, Parkinson’s, Huntington’s and multiple sclerosis (MS). Thus, BACH1 inhibitors, by inducing *HMOX1*, could be developed as neuroprotective agents.

Despite their therapeutic potential, so far only very few BACH1 inhibitors have been reported. The compound most widely used to inhibit BACH1 is the oxidized form of its endogenous ligand heme. Hemin binds to BACH1, impairing its binding to DNA, promoting its nuclear export and its degradation via the E3 ligase Fbxo22 [17] [18] [19]. Hemin is a very potent inhibitor, but its clinical use is limited by the toxic nature of free heme [20]. Other BACH1 inhibitor is the natural non-psychotropic phytocannabinoid cannabidiol (CBD). We have shown that CBD targets BACH1 for degradation and, consequently, induces *HMOX1* [9]. CBD is neuroprotective in a murine MS model [21] and has been tested in various animal and cell models of other NDs [22], and some of its beneficial effects could indeed be related to its inhibitory effect on BACH1. Another reported BACH1 inhibitor is the synthetic compound HPP-4382 [23] that, to the best of our knowledge, has not yet been tested in any disease models.

Pharmacological activation of NRF2 is also beneficial in most pathologies with underlying oxidative stress and inflammation [24]. Most compounds that stabilise NRF2 are either electrophilic molecules that covalently modify certain cysteine residues in KEAP1, impairing its activity [25], or inhibitors that interfere with the binding between NRF2 and KEAP1 [26]. Stabilization of NRF2 triggers the transcription of multiple enzymes involved in scavenging ROS and in generating antioxidant molecules like glutathione [27] [28] while leading to a reduction of pro-inflammatory cytokines such as IL-6 [29] [30]. Thus NRF2 activators have also been studied for their potential neuroprotective effect [31]. In fact, the activation of the NRF2 pathway has been shown to be neuroprotective in various models of NDs such as MS and Huntington disease (HD) [32] [33]. HD is caused by a mutation in the huntingtin (HTT) gene; this mutant HTT tends to form aggregates that lead to the dysfunction and eventual death of neurons within the striatum [34]. HD presents high levels of oxidative stress, brain inflammation and impaired NRF2 activity [35], and NRF2 activation has been shown to repress proinflammatory processes in preclinical HD models [35]. Furthermore, treatment with the NRF2 activator dimethyl fumarate (DMF) resulted in improved motor function in a mouse HD model [36].

While NRF2 activators induce the expression of many cytoprotective genes, BACH1 inhibitors activate only a few, although they are extremely potent at inducing *HMOX1*. Thus, the individual knockdown of KEAP1 (leading to constitutive NRF2 activation) or BACH1 in HaCaT cells induced the expression of *HMOX1* by 2.3- and 136-fold, respectively, whereas the combined knockdown of KEAP1 and BACH1 resulted in a 388-fold increase of *HMOX1* mRNA [37]. Consequently, complementing NRF2 activation with BACH1 inhibition would result in a highly robust anti-inflammatory and antioxidant response, and potentially a better therapeutic effect.

Cannabidiol has sparked attention due to its therapeutic potential, and there is considerable interest in synthesizing new CBD derivatives with improved pharmacological and clinical properties. In this study, we report the identification of a novel CBD derivative that is highly effective at both inhibiting BACH1 and activating NRF2, and show its cytoprotective efficacy in cell models of relevance to Huntington’s disease.

## MATERIALS AND METHODS

### Cell culture

The following cell lines were used in this study: HaCaT, THP1, SH-SY5Y, Hepa 1c1c7, A549 and Q7/Q111. HaCaT cells have been validated by STR *profiling* and were routinely tested for mycoplasma. HaCaT-ARE-Luc cells were generated as described previously [38]. HaCaT NRF2-KO cells were produced using the CRISPR-Cas9 system as described [9]. CRISPR-edited HaCaT BACH1-KO and HaCaT NRF2-KO/BACH1-KO cells were generated by transfecting either HaCaT WT or HaCaT NRF2-KO cells with two different pLentiCRISPR-v2 (a gift from Dr Feng Zhang, Addgene plasmid #52961) containing each one a guide RNA against the first exon and the second exon of BACH1, respectively (5’-CGATGTCACCATCTTTGTGG-3’, 5’-GACTCTGAGACGGACACCGA-3’). All CRISPR-edited cell lines were selected for two days using puromycin, cells were clonally selected by serial dilution and positive clones were identified, as previously described [39]. Control cells, referred to as HaCaT wild type (HaCaT WT), are the pooled population of surviving cells transfected with an empty pLentiCRISPRv2 vector treated with puromycin. THP1, SH-SY5Y, Hepa-1c1c7 and A549 cells were obtained from ATCC and were routinely tested for mycoplasma. Clonal striatal cell lines established from E14 striatal primordia of Hdh^Q111/Q111^ (mutant) and Hdh^Q7/Q7^ (wild-type) knock-in mouse littermates and immortalized as previously described [40]. Cell lines were grown in RPMI (HaCaT, THP1), DMEM (A549), DMEM/F12 (SH-SY5Y) or alpha-MEM (Hepa-1c1c7) with 10% FBS at 37 °C and 5% CO_2_. Striatal Q7 and Q111 cells were maintained in DMEM containing 25⍰1mM D-glucose, 11⍰mM L-glutamine, 10% fetal bovine serum (FBS), 1⍰mM sodium pyruvate, and 400⍰μg/mL Geneticin (Invitrogen, Carlsbad, CA) and were incubated at 33°C with 5% CO2.

### Antibodies and Reagents

Antibodies recognizing Beta-Actin (C-4) and anti-BACH1 (F-9) were obtained from Santa Cruz Biotechnology (Dallas, Texas, USA). anti-NRF2 (D1Z9C) was obtained from Cell Signalling Technology (Danvers, MA, USA) and anti-HMOX1 was purchased from Biovision (San Francisco, CA, USA). The siRNAs used as control or against KEAP1 were the SMART pool: ON-Target Plus from Dharmacon (Lafayette, CO, USA). CBD (Fig. 1B (1)) was isolated from the Cannabis strain Carmagnola [41].

**Fig. 1.**
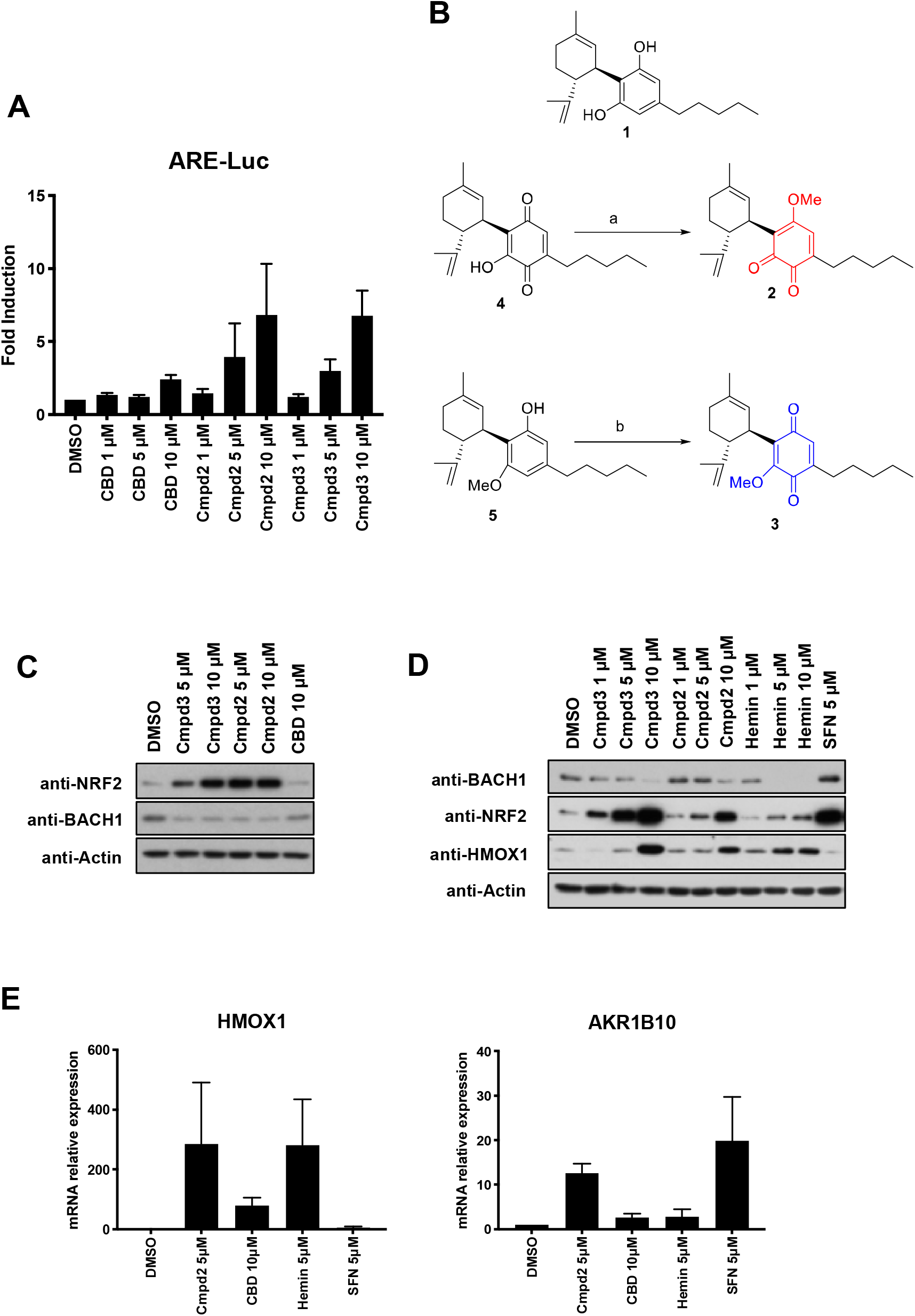
Screening of CBD derivatives and characterization of compound 2 as a BACH1 inhibitor and NRF2 inducer. **A)** HaCaT-ARE-Luc cells were treated with either DMSO or increasing concentrations of CBD, compound 2 (Cmpd2) or compound 3 (Cmpd3) for 6lll h. Luciferase activity was measured in the cell lysates and expressed as RLU (× 10^4^). Data represent means ± SD (n = 4) and are expressed relative to untreated cells. **B)** Schematic representation of the sinthesis of the isomeric methoxyquinones 2 (compound 2) and 3 (compound 3) from 1 (CBD). (a): MeI, NaHCO3, DMF, 49%; (b) SIBX, EtOAc, 51%. **C)** HaCaT cells were incubated with either DMSO, compound 3 (Cmpd3), compound 2 (Cmpd2) or CBD. Three hours later, cells were lysed and samples were analysed by Western Blot. **D)** HaCaT cells were incubated with either DMSO, or increasing concentrations of compound 3 (Cmpd3), compound 2 (Cmpd2), hemin or sulforaphane (SFN). Three hours later, cells were lysed and levels of BACH1, NRF2, *HMOX1* and Actin were analysed by Western Blot. A representative blot is shown; the corresponding quantifications of BACH1, NRF2 and *HMOX1* protein levels are shown in Suppl. Fig. S1A. **E)** HaCaT cells were treated with either DMSO, compound 2 (5 μM), CBD (10 μM), Hemin (5 μM) or SFN (5 μM) for 8 hours. The mRNA levels of *HMOX1* and *AKR1B10* were quantified by real-time PCR and the data were normalised using HPRT1 as an internal control. Data represent means ± SD (n = 3) and are expressed relative to the DMSO sample. * P≤0.05, **P≤ 0.01, ***P ≤ 0.001.

### Synthesis of the CBD derivatives

#### *O*-Methyl *ortho*-cannabinodiolquinone (Compound 2)

To a cooled (ice bath) solution of cannabidiolquinone (Fig. 1B (4)) (1.87 g, 5.69 mmol) [42] in DMF (19 mL), solid NaHCO_3_ (0.95 g, 11.31 mmol) was added. After stirring for 10 min., methyl iodide (1.75 g, 28.11 mmol) was added, and the resulting mixture was stirred overnight at room temperature. The reaction was worked up by dilution with water (75 mL) and EtOAc (75 mL), and, after a 5 minute stirring, the two layers were separated. The lower organic layer was washed with water (2 × 25 mL), dried (Na_2_SO_4_) and evaporated to obtain a dark oily residue, that was purified by gravity column chromatography on silica gel, using an EtOAc gradient in hexane (from 1:99 to 4:96). 960 mg (49%) *O*-methyl ortho-cannabinodiolquinone were obtained as a red oil. ^1^H-NMR (300 MHz, CDCl_3_) δ = 6.37 (t, J=1.5, 1H), 5.10 (s, 1H), 4.59 – 4.48 (m, 2H), 3.87 (s, 3H), 3.82-3.67 (m, 1H), 2.70 (td, J=11.1, 10.7, 3.2, 1H), 2.36 (ddd, J=10.2, 6.2, 1.5, 2H), 2.29 – 2.10 (m, 1H), 2.07 – 1.91 (m, 1H), 1.81 – 1.59 (m, 3H), 1.69 (s, 3H), 1.64 (s, 3H), 1.57 – 1.42 (m, 2H), 1.42 – 1.24 (m, 4H), 0.97 – 0.83 (m, 3H).

#### *O*-Methyl cannabidiolquinone (Compound 3)

To a solution of *O*-methyl cannabidiol (Fig. 1B (5)) [43] (2.25 g, 6.85 mmol) in EtOAc (150 mL), SIBX (42.7%, 13.5 g, 20.59 mmol) was added, and the mixture was stirred at room temperature for 24 h. The reaction was worked up by filtration, and he filtrate was washed with sat. NaHCO_3_ (3 × 100 mL), dried (Na_2_SO_4_), and evaporated. A dark red oil was obtained, that was purified by gravitiy colum chromatography (silica gel, EtOAc/hexanes 4:96 as eluant) to afford 1.19 g (51%) of *O*-methyl *ortho*-cannabidiolquinone as a dark red oil. ^1^H-NMR (300 MHz, CDCl_3_) δ = 6.84 (s, 1H), 5.08 (s, 1H), 4.61 – 4.51 (m, 2H), 3.89 (s, 3H), 3.74 – 3.61 (m, 1H), 2.76 – 2.62 (m, 1H), 2.35 (t, J=7.7, 2H), 2.27-2.10 (m, 1H), 2.03 – 1.91 (m, 1H), 1.80-1.59 (m, 3H), 1.66 (s, 3H), 1.63 (s, 3H), 1.49 (m, 2H), 1.39 – 1.24 (m, 4H), 0.95-0.82 (m, 3H).

### Quantitative real time PCR (rt-qPCR)

Samples were lysated and homogenised using the QIAshredder (Qiagen, Hilden, Germany) and RNA was extracted using RNeasy kit (Qiagen). 500 ng of RNA per sample was reverse-transcribed to cDNA using Omniscript RT kit (Qiagen) supplemented with RNase inhibitor (Invitrogen, Waltham, MA, USA) according to the manufacturer’s instructions. Resulting cDNA was analysed using TaqMan Universal Master Mix II (Life Technologies, Carlsbad, CA, USA). Gene expression was determined by the comparative ΔΔCT method, using a QuantStudio 7 Flex qPCR machine. All experiments were performed at least in triplicates and data were normalized to the housekeeping gene HPRT1. The TaqMan probes used are:

HPRT1 Hs02800695_m1 HPRT1TaqMan(s)
HMOX1 Hs01110250_m1 *HMOX1* TaqMan(s)
AKR1B10 Hs00252524_m1 AKR1B10 TaqMan (s)
NQO1 Hs01045993_g1 NQO1 TaqMan (m)

### Cell lysis and western blot protocol

Cells were washed and harvested in ice-cold phosphate-buffered saline (PBS) and lysed in RIPA buffer supplemented with phosphate and protease inhibitors [50 mM Tris-HCl pH 7.5, 150mM NaCl, 2mM EDTA, 1% NP40, 0.5% sodium deoxycholate, 0.5 mM Na3VO4, 50 mM NaF, 2 μg/ml leupeptine, 2 μg/ ml aprotinin, 0.05 mM pefabloc]. Lysates were sonicated for 15 s at 20% amplitude and then cleared by centrifugation for 15 min at 4 °C. Protein concentration was established using the BCA assay (Thermo Fisher Scientific, Waltham, MA, USA). Lysate was mixed with SDS sample buffer and boiled for 7 min at 95 °C. Equal amounts of protein were separated by SDS-PAGE, followed by semidry blotting to a polyvinylidene difluoride membrane (Thermo Fisher Scientific). After blocking of the membrane with 5% (w/v) TBST non-fat dry milk, primary antibodies were added overnight. Appropriate secondary antibodies coupled to horseradish peroxidase were detected by enhanced chemiluminescence using ClarityTM Western ECL Blotting Substrate (Bio-Rad, Hercules, CA, USA). Resulting protein bands were quantified and normalised to each lane’s loading control using the ImageStudio software (LI-COR).

### siRNA cell transfections

The day before the transfection cells were plated into 6-well plates to 70-90% confluency. The siRNA and Lipofectamine RNAiMAX (Invitrogen, Carlsbad, CA, USA) were individually diluted in Opti-MEM (Life Technologies) and incubated for 10 min at room temperature. Diluted siRNA was added to the diluted Lipofectamine solution (1:1 ratio) and further incubated for 15 min. The complex was added to the cells and incubated in a humidified incubator at 37 °C and 5% CO2 for 36 h prior treatment and lysis.

### Cell viability assays

To assess cell viability of HaCaT cells after drug treatment we performed an Alamar Blue assay. One day before the experiment, HaCaT cells were seeded into a 96-well plate to 50-60% confluency. Cells were treated with the corresponding compounds for 16 hours and Alamar Blue (Thermo Fisher Scientific) was added to the wells. After 4 hours of incubation at 37 °C the fluorescence was measured using a microplate reader (Spectramax m2) and viability was calculated relative to the DMSO treated control. For the in vitro HD model we used ST*Hdh*^Q7/Q7^ and ST*Hdh*^Q111/Q111^ cells, which express either a wild type or a mutated form of the huntingtin protein [44]. These cells were cultured at 33 °C and 5% CO_2_ in DMEM supplemented with 10 % FBS, 2 mM L-glutamine and 1 % (v/v) penicillin/streptomycin (Epub 2000/11/25). ST*Hdh*^Q7/Q7^ and ST*Hdh*^Q111/Q111^ cells (10^4^ cells/well) were seeded in DMEM supplemented with 10 % FBS in 96 well plates incubated with increased concentrations of compound 2 and treated with 3-NP at 10 mM (#N5636, Sigma). Then, 3-NP-induced cytotoxicity was measured by fluorescence using the die YOYO-1 (#Y3601, Life Technologies). Treated cells were placed in an Incucyte FLR imaging system and the YOYO-1 fluorescence was measured after several time points. Object counting analysis is performed using the Incucyte FLR software to calculate the total number of YOYO-1 fluorescence positive cells and total DNA containing objects (endpoint). The cytotoxicity index is calculated by dividing the number of YOYO-1 fluorescence positive objects by the total number of DNA containing objects for each treatment group.

### Luciferase assays

HaCaT-ARE-Luc cells were stimulated with either Sulforaphane (SFN) (5 μM) or with increasing concentrations of cannabinoids for 6 h. After the treatment the cells were washed twice in PBS and lysed in 25 mM Tris-phosphate pH 7.8, 8 mM MgCl_2_, 1 mM DTT, 1% Triton X-100, and 7% glycerol during 15 min at room temperature in a horizontal shaker. After centrifugation, luciferase activity in the supernatant was measured using a GloMax 96 microplate luminometer (Promega) following the instructions of the luciferase assay kit (Promega, Madison, WI, USA). Results are expressed in RLU over control untreated cells.

### ROS determination

The intracellular accumulation of ROS was detected using 2’,7-’dihydrofluorescein-diacetate (DCFH-DA). HaCaT cells (15×10^3^ cells/well) were cultured in a 96-well plate in DMEM supplemented with 10% FBS until cells reached 80% confluence. Cells were pre-treated with the compounds for 30 min and treated with 0,4 mM Tert-butyl-hydroperoxide (TBHP). At the indicated time points, the cells were incubated with 10 μM DCFH-DA in the culture medium at 37°C for 30 min. Then, the cells were washed with PBS at 37°C and the production of intracellular ROS was measured by DCF fluorescence and detected using the Incucyte FLR software. The data were analyzed by the total green object integrated intensity (GCUxμm^2^xWell) of the imaging system IncuCyte HD (Sartorius, Göttingen, Germany).

### NQO1 activity assay

Cells were lysed in digitonin (0.8 g/L in 2 mM EDTA, pH 7.8), and the lysates were subjected to centrifugation at 4°C (15,000 × g for 10 minutes). The enzyme activity of NQO1 was measured in lysate supernatants using menadione as a substrate as previously described [45].

### Statistical analysis

Most experiments were repeated 3–5 times with multiple technical replicates to be eligible for the indicated statistical analyses. Data were analysed using Graphpad Prism statistical package. All results are presented as mean ± SD unless otherwise mentioned. When applicable, the differences between groups were determined by either one-way ANOVA or 2-way ANOVA. A P value of < 0.05 was considered significant. *P ≤ 0.05, **P ≤ 0.01, ***P ≤ 0.001.

## RESULTS AND DISCUSSION

### Identification of CBD derivatives that activate ARE-dependent gene expression

We have recently shown that CBD is a potent BACH1 inhibitor, but a very weak NRF2 activator in HaCaT cells [9]. In search for CBD derivatives with higher potency in activating NRF2, we screened a library of semi-synthetic cannabidiolquinone derivatives in the NRF2 reporter cell line HaCaT ARE-Luc, identifying the isomeric quinoids O-methyl ortho-cannabidiolquinone (hereinafter referred to as compound 2) and O-methyl cannabidiolquinone (hereinafter referred to as compound 3) as reporter activators in a concentration-dependent manner (Fig. 1A).

Compared to CBD, compounds 2 and 3 are characterized by the oxidation of the resorcinyl core to a methoxylated quinone moiety (Fig. 1B). Early studies preceding the discovery of NRF2 had shown that among diphenols, only 1,2 diphenols (catechols) and 1,4-diphenols (hydroquinones), but not 1,3-diphenols (resorcinols), were inducers of the cytoprotective enzyme NAD(P)H:quinone oxidoreductase 1 (NQO1) [46] which is now widely accepted as a classical NRF2-regulated protein [47]. This realization established that oxidative lability among diphenols, i.e. their ability to form quinones in the presence of oxygen, was essential for inducer activity. This notion was further strengthened by the finding of a linear correlation between the NQO1 inducer activity and the ability to release an electron among inducers of different classes, including diphenols, phenylpropenoids, and flavonoids [48]. Subsequent work showed that the stronger the electron-attracting property of a quinone, the greater its inducing potency [49], implicating oxidation of cysteine sensors of KEAP1 by quinones in the mechanism of NRF2 activation. Experiments using recombinant KEAP1 in the presence or absence of Cu^++^ and oxygen firmly established that oxidizable diphenols are not NRF2 activators themselves, but their corresponding quinones are the ultimate inducers [50].

Diphenols have been proposed as pro-drugs to combat oxidative stress based on the concept that such compounds are chemically converted to their active quinone forms by the oxidative environment that they are designed to protect against [51]. It is therefore highly likely that the NRF2 inducer property of these compounds is conferred by their quinone functionalities.

To validate the results from the screen, we first compared the effect of compound 2 and 3 to that of CBD on the protein levels of BACH1 and NRF2 in HaCaT cells. Both compounds were more potent than CBD at stabilising NRF2, fully retaining the ability to reduce the levels of BACH1 (Fig. 1C).

### Validation of compound 2 as a dual BACH1 inhibitor and NRF2 inducer

To further characterise the effect of compound 2 and compound 3 on BACH1 and NRF2, we compared them with the BACH1 inhibitor hemin and the NRF2 activator sulforaphane (SFN) (Fig. 1D and quantification in Suppl. Fig S1A). Both compound 2 and compound 3 reduced BACH1 protein levels. Consistent with the negative regulation of *HMOX1* expression by BACH1, treatment with compound 2 or compound 3 increased the protein levels of *HMOX1*. When compared with hemin, both compounds at a concentration of 10 μM showed similar efficacy in decreasing BACH1 levels as that of hemin at a concentration of 1 μM. Furthermore, both compound 2 and compound 3 stabilised NRF2 in a concentration-dependent manner, and this effect was comparable to the effect of SFN. As compound 3 was more toxic (Suppl. Fig. S1B) and less stable at room temperature (Suppl Fig. S1C), we focused on compound 2 for the rest of this study.

To confirm that the activation of NRF2 by compound 2 was not limited to HaCaT cells, we tested this compound in Hepa1c1c7 cells, a cell line widely used to study NRF2 activation [52]. Our results showed that in Hepa1c1c7 cells, compound 2 stabilised NRF2 and induced *HMOX1* (Suppl. Fig. S1D). Compound 2 also dose-dependently induced the enzyme activity of the classical NRF2 target, NAD(P)H:quinone oxidoreductase 1 (NQO1) (Suppl. Fig S1E).

We have previously shown that in HaCaT cells, *HMOX1* expression is an excellent surrogate for BACH1 activity as it is regulated primarily by BACH1 and not by NRF2, while *AKR1B10* is a surrogate for NRF2 activity and does not respond to BACH1 modulation [9] (also see Suppl. Fig. S2). To further validate compound 2 as a dual BACH1 inhibitor and an NRF2 inducer, we analysed its effect on the expression of these surrogate markers in HaCaT cells. As shown in Fig. 1E, compound 2 was more potent than CBD at inducing *HMOX1* and *AKR1B10* expression. As expected, SFN induced the expression of *AKR1B10*, but not *HMOX1*, whereas hemin induced *HMOX1*, but not *AKR1B10* expression.

### Characterisation of the effect of compound 2 on NRF2 and BACH1

To further characterise the effect of compound 2, we tested whether in a similar way to CBD, the compound 2-mediated induction of *HMOX1* was dependent on BACH1. As observed in Fig 2A, the ability of both compound 2 and hemin to induce *HMOX1* was abolished in BACH1-KO cells (validation of BACH1-KO cells in Suppl Fig S2A and Suppl S2B). On the other hand, BACH1 knockout did not impair the effect of compound 2 on *AKR1B10* (Fig. 2B) or NRF2 (Fig. 2C), showing that NRF2 activation by compound 2 does not depend on BACH1. Similarly, the inhibition of BACH1 by compound 2 was still evident in NRF2-KO cells (Fig. 2D), indicating that the two effects of compound 2 are independent (validation of NRF2-KO cells in Suppl Fig. S2C and Suppl Fig. S2D).

**Fig. 2.**
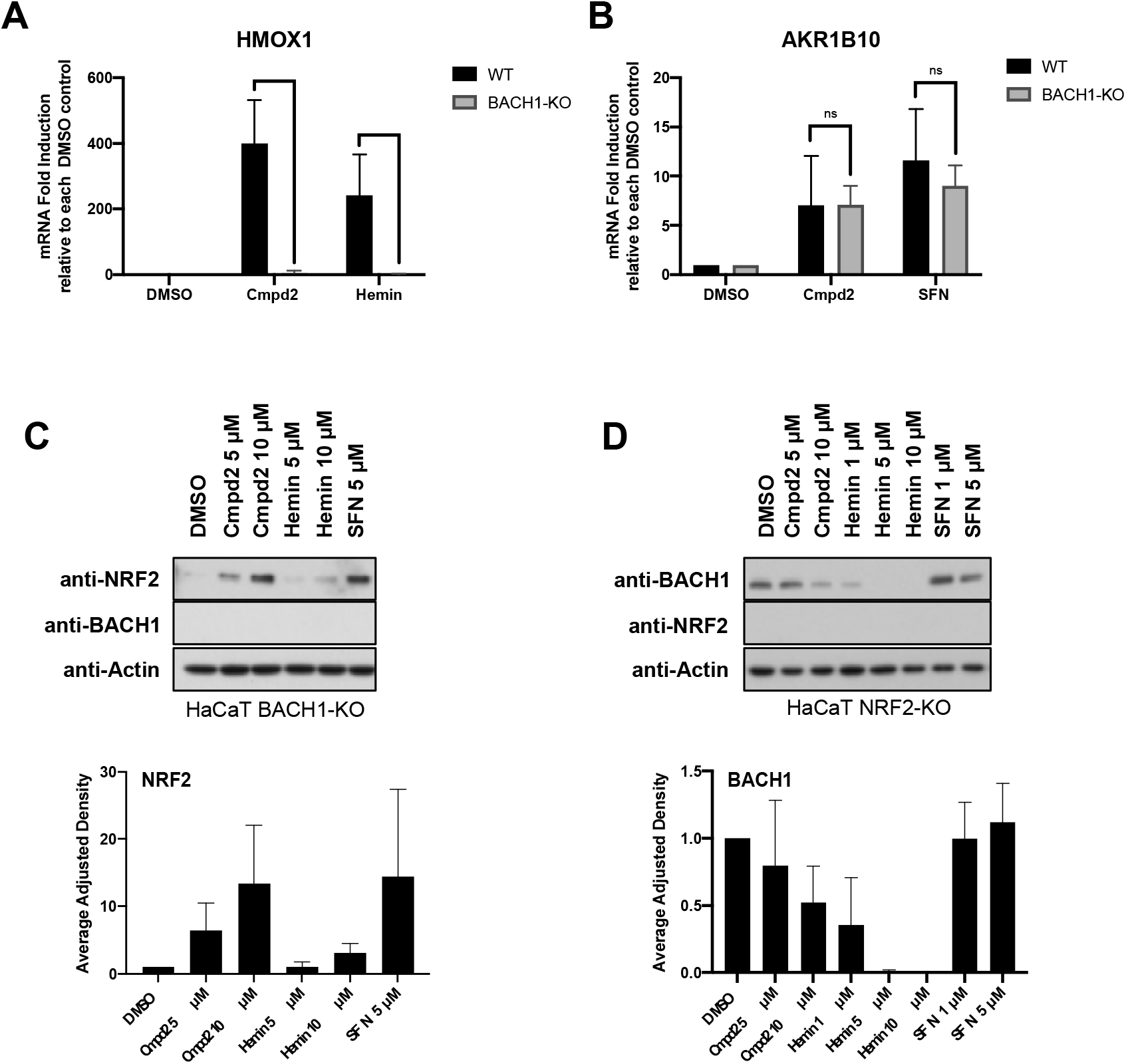
Compound 2-mediated NRF2 induction and BACH1 inhibition are not interdependent. **A)** HaCaT WT and BACH1-KO cells were treated for 8 hours with DMSO, compound 2 (10 μM) or Hemin (10 μM). Cells were lysed and *HMOX1* mRNA levels were quantified (n=3). To compare the *HMOX1* induction upon compound 2 or Hemin treatment in each cell line, the levels of *HMOX1* in either treated WT or treated BACH1-KO samples were compared against the basal level of *HMOX1* in DMSO WT or DMSO BACH1-KO respectively (*HMOX1* levels in DMSO samples were set in both cases as 1). **P≤ 0.01, ***P ≤ 0.001. **B)** HaCaT WT and BACH1-KO cells were treated as indicated in A. Cells were lysed and *AKR1B10* mRNA levels were quantified by real-time PCR and the data were normalised using HPRT1 as an internal control (n=3). As described in A, *AKR1B10* levels in DMSO WT and BACH1-KO samples were set in both cases as 1. **C)** HaCaT BACH1-KO cells were incubated with either DMSO or different concentrations of compound 2, Hemin or SFN. After 3 hours, cells were harvested and lysates were analysed by Western Blot. Upper panel is a representative western blot and the bottom panel shows the quantification of NRF2 protein levels against the loading control actin. Data represent means ± SD (n = 3) and are expressed relative to the DMSO treated samples. **D)** HaCaT NRF2-KO cells were treated and harvested as described in C and samples were analysed by Western Blot. Upper panel is a representative western blot and the bottom panel shows the quantification of BACH1 protein levels against the loading control actin. Data represent means ± SD (n = 3) and are expressed relative to the DMSO samples.

Next, we compared the kinetics of *HMOX1* and *AKR1B10* expression in response to compound 2, hemin and SFN in WT and NRF2-KO cells (Fig 3A, 3B and 3C). Compound 2 increased *HMOX1* mRNA levels in an NRF2-independent manner, with similar kinetics as hemin; starting as early as two hours and reaching its peak around eight hours (Fig. 3A and 3C). As previously observed, SFN induced *HMOX1* very weakly at the 8-hour time point (Fig. 3B). On the other hand, compound 2 induced *AKR1B10* expression in an NRF2-dependent manner with similar kinetics as SFN, starting around four hours and reaching its peak around eight hours (Fig. 3A and 3B). As expected, hemin did not induce *AKR1B10* (Fig. 3C). These results further confirm that compound 2 has dual activity targeting both BACH1 and NRF2.

**Fig. 3.**
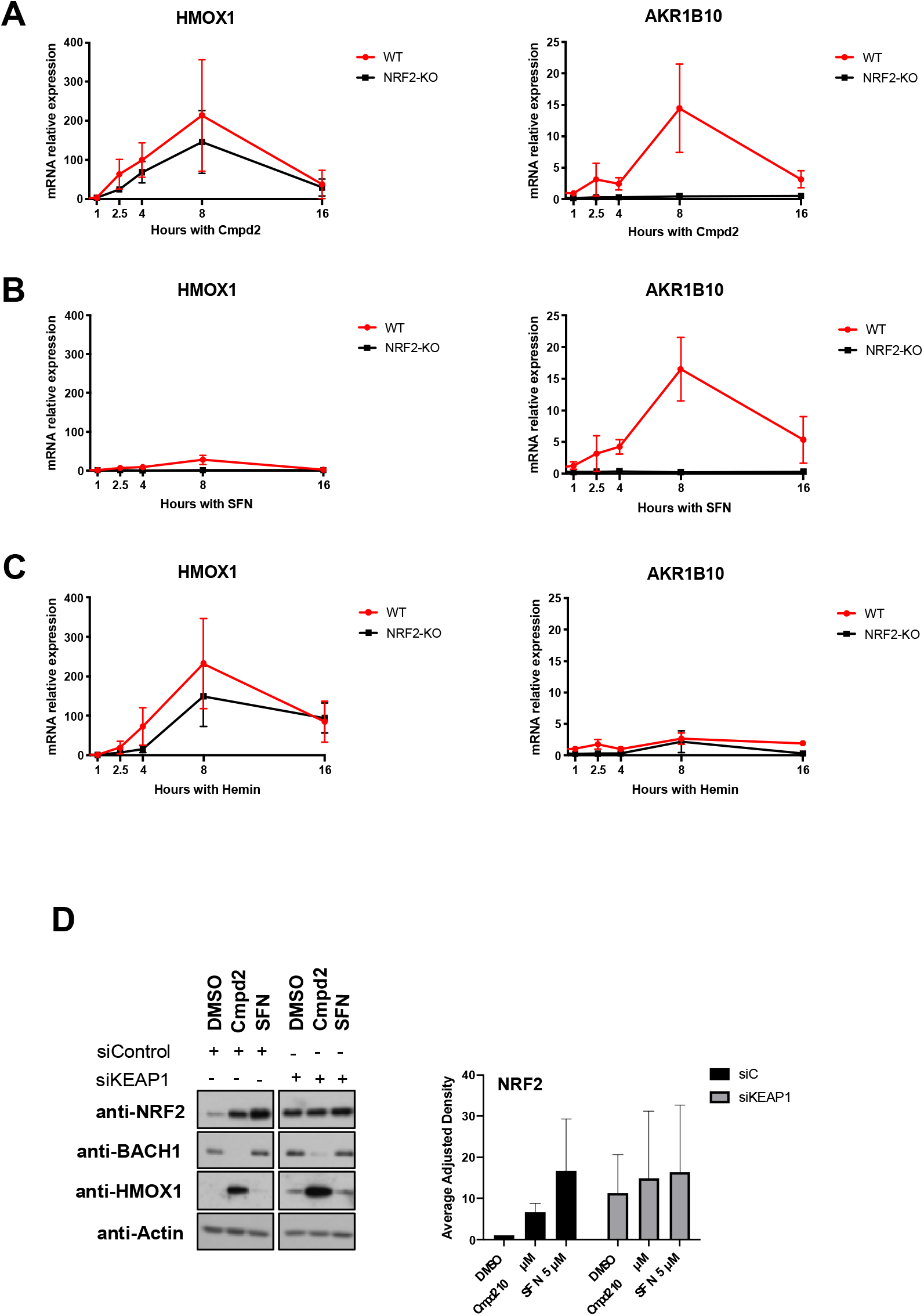
Compound 2 inhibits BACH1 in a NRF2 and KEAP1 independent manner, and stabilises NRF2 in a KEAP1-dependent manner. **A)** HaCaT WT and NRF2-KO cells were treated with compound 2 (10 μM) and were harvested after 1, 2:30, 4, 8 or 16 hours. Samples were lysed and *HMOX1* (left panel) or *AKR1B10* (right panel) mRNA levels were quantified by real-time PCR using HPRT1 as an internal control. *HMOX1* and *AKR1B10* levels in DMSO samples were set in both cell lines as 1 with the purpose of comparing the induction after compound 2 treatment in WT and NRF2-KO cells. Data represent means ± SD (n = 3). *P ≤ 0.05, **P≤ 0.01 **B)** HaCaT WT and NRF2-KO cells were treated with SFN (5 μM) for 1, 2:30, 4, 8 or 16 hours. mRNA *HMOX1* and *AKR1B10* levels were analysed as described in A (n=3). **C)** HaCaT WT and NRF2-KO cells were treated with SFN (5 μM) for 1, 2:30, 4, 8 or 16 hours. mRNA *HMOX1* and *AKR1B10* levels were analysed as described in A (n=3). **D)** HaCaT cells were transfected with either siControl or siKEAP1. 36 hours later cells were treated with either DMSO, compound 2 (10 μM) or SFN (5 μM) for 3 hours. Cells were lysed and samples were analysed by Western Blot. Left panel is a representative western blot and right panel shows the quantification of NRF2 protein levels against the loading control. Data represent means ± SD (n = 2) and are expressed relative to the siControl DMSO sample; the corresponding quantifications of BACH1 and *HMOX1* protein levels are shown in Suppl. Fig. S3A

Based on the strong effect of compound 2 increasing NRF2 levels, we hypothesise that this stabilisation must be a consequence of impairing the activity of its main regulator KEAP1. To test this possibility, we compared the effect of compound 2 on NRF2 levels in HaCaT cells treated with either siRNA control or an siRNA against KEAP1. The stabilisation of NRF2 induced by compound 2 was clearly impaired in KEAP1-silenced cells (Fig. 3D and Suppl Fig. S3A). We also tested compound 2 and in the lung cancer cell line A549 bearing a loss-of-function mutation in KEAP1, and found that the NRF2 levels were not significantly changed by compound 2 (Suppl. Fig S3B). Collectively, these results suggest that NRF2 activation by compound 2 requires KEAP1, and is independent of its effect on BACH1.

### Compound 2 protects against oxidative stress in cell-based models

The characterization of the effect of compound 2 on BACH1 and NRF2 was performed in HaCaT cells because this is a validated model where we can individually assess the compound activity on BACH1 and NRF2. However, to answer whether compound 2 could have a therapeutic benefit in NDs, we used two cell lines of relevance to Huntington’s disease as models for inflammation and oxidative stress in the brain. First, we tested for target engagement in macrophage-like THP1 cells and the neuroblastoma cell line SH-SY-5Y. Compound 2 treatment efficiently decreased the levels of BACH1, induced *HMOX1* and stabilised NRF2 in these cell lines (Fig. 4A). As ROS play a role in HD, and may be the initial molecules leading to apoptosis, we compared the ability of compound 2 to decrease the levels of ROS generated by treatment with tert-butyl hydroperoxide (tBHP) with that of SFN and dimethyl fumarate (DMF) at different time points. DMF is a NRF2 inducer that has been approved for the treatment of multiple sclerosis by the FDA. At all time points tested, compound 2 decreased the ROS levels more efficiently than either SFN or DMF (Fig. 4B).

**Fig. 4.**
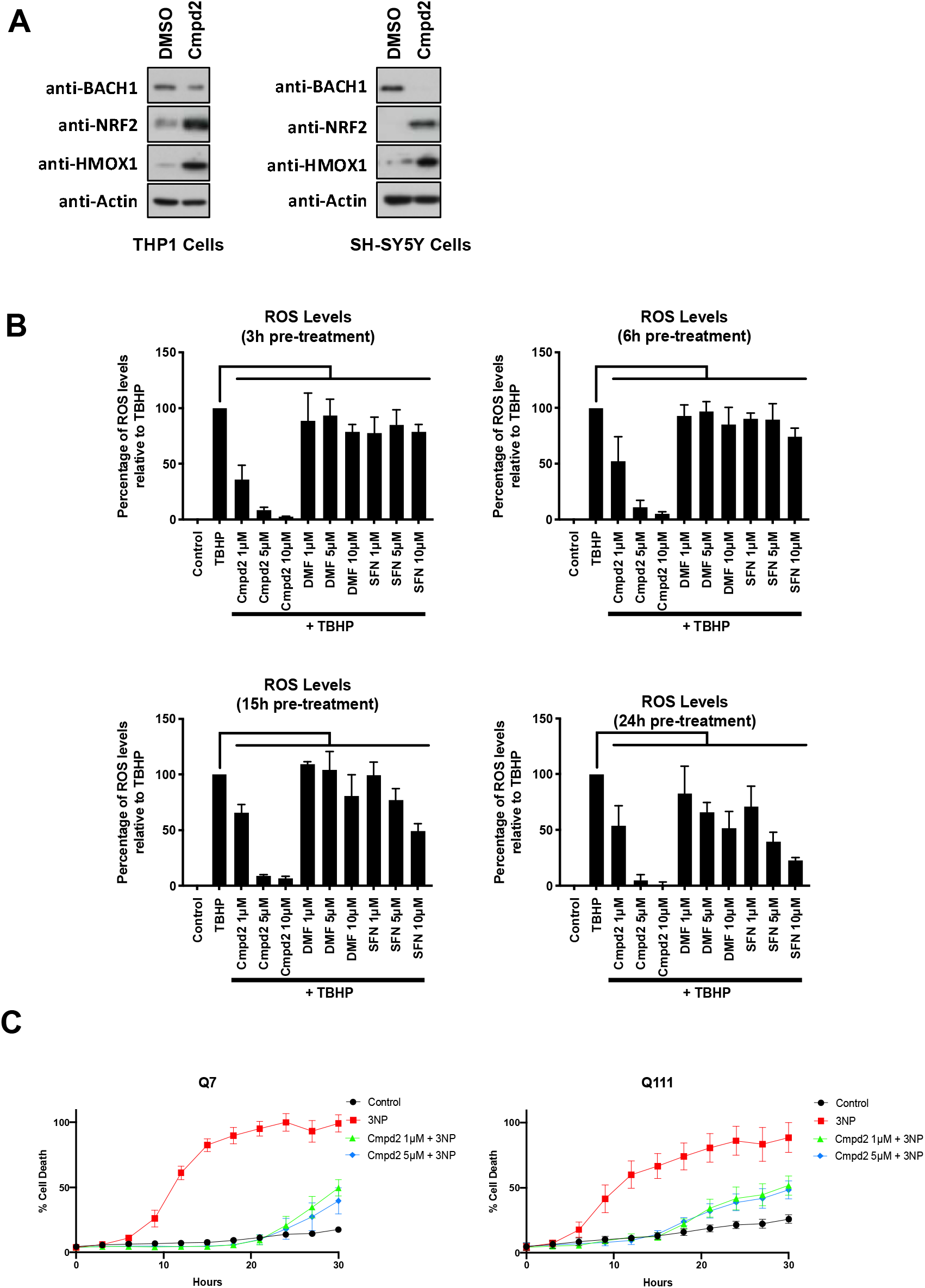
Compound 2 stabilizes NRF2 and induces *HMOX1* in macrophages and neural cells and protects cells from 3-NP mediated cytotoxicity in an *in vitro* HD model. **A)** THP1 or SH-SY5Y cells were treated with either DMSO or compound 2 (10 μM). After 3 hours, cells were lysed and BACH1, NRF2, *HMOX1* and Actin protein levels were analysed by Western Blot. **B)** HaCaT cells were treated as indicated for 3h, 6h, 15h or 24h and the detection and quantification of ROS (DCF fluorescence) was measured by fluorescence microscopy. Data represent means ± SD (n = 3) and are expressed relative to control cells. *P ≤ 0.05, **P≤ 0.01, ***P ≤ 0.001. **C)** Striatal Q7 and Q111 cells were pre-treated with either DMSO or increasing concentrations of compound 2 for 6⍰h and then exposed to 10⍰mM 3-NP in low glucose media for an additional 24⍰h. Cell death was monitored using the INCUCYTE. ***P ≤ 0.001.

Furthermore, we tested whether compound 2 could provide neuroprotection in an *in vitro* model for HD, using conditionally-immortalized striatal neuronal progenitor cell lines expressing endogenous levels of normal (STHdhQ7/Q7) and mutant huntingtin (STHdhQ111/Q111). 3-nitropropionic acid (3-NP) is a natural toxin that significantly induces oxidative damage in the brain, producing striatal lesions that are very similar to those of patients with HD [53]; and thus, it is widely used as the model agent to mimick HD. When used in the striatal neural cell lines, compound 2 treatment significantly reduced the cytotoxicity caused by 3-NP in both cell lines (Fig. 4C), confirming its protective effect.

Our data demonstrate that O-methyl ortho-cannabidiolquinone (compound 2) has dual activity as a BACH1 inhibitor and an NRF2 activator. This is a very attractive profile for drugs against oxidative stress and/or inflammatory associated conditions, such as most neurodegenerative disorders, and our data on a HD cell model provide a solid rationale for future studies *in vivo* using NDs models. It is noteworthy that with aging, NRF2 signaling is impaired [54] [55] [56]. A comparison of NRF2 activation by sulforaphane in bronchial epithelial cells isolated from young human subjects (21-29 years) and older (60-69 years) individuals, non-smokers, has shown that the inducibility of the NRF2-mediated cytoprotective responses is diminished in cells from older adults [57]. Importantly, this is accompanied by an increased expression of BACH1 [57]. Conversely, silencing of BACH1 enhances sulforaphane-induced expression of NRF2-target genes in cells from older subjects [58]. Thus, our findings suggest that compounds such as compound 2 may have beneficial effects in aging.

## AUTHORS CONTRIBUTION

LC, JU, MP and DC conducted the experiments and were responsible for initial data analysis, figure preparation and statistical analysis. GA provided resources. LV and EM and ADK had a leading contribution in the design of the study, and an active role in the discussion and interpretation of the whole dataset. LV wrote the original draft of the manuscript. LV, EM, ADK, GA and LC, reviewed and edited the manuscript. Funding acquisition LV and EM. All the authors take full responsibility for the work.

---

**Suppl. Figure S1.**
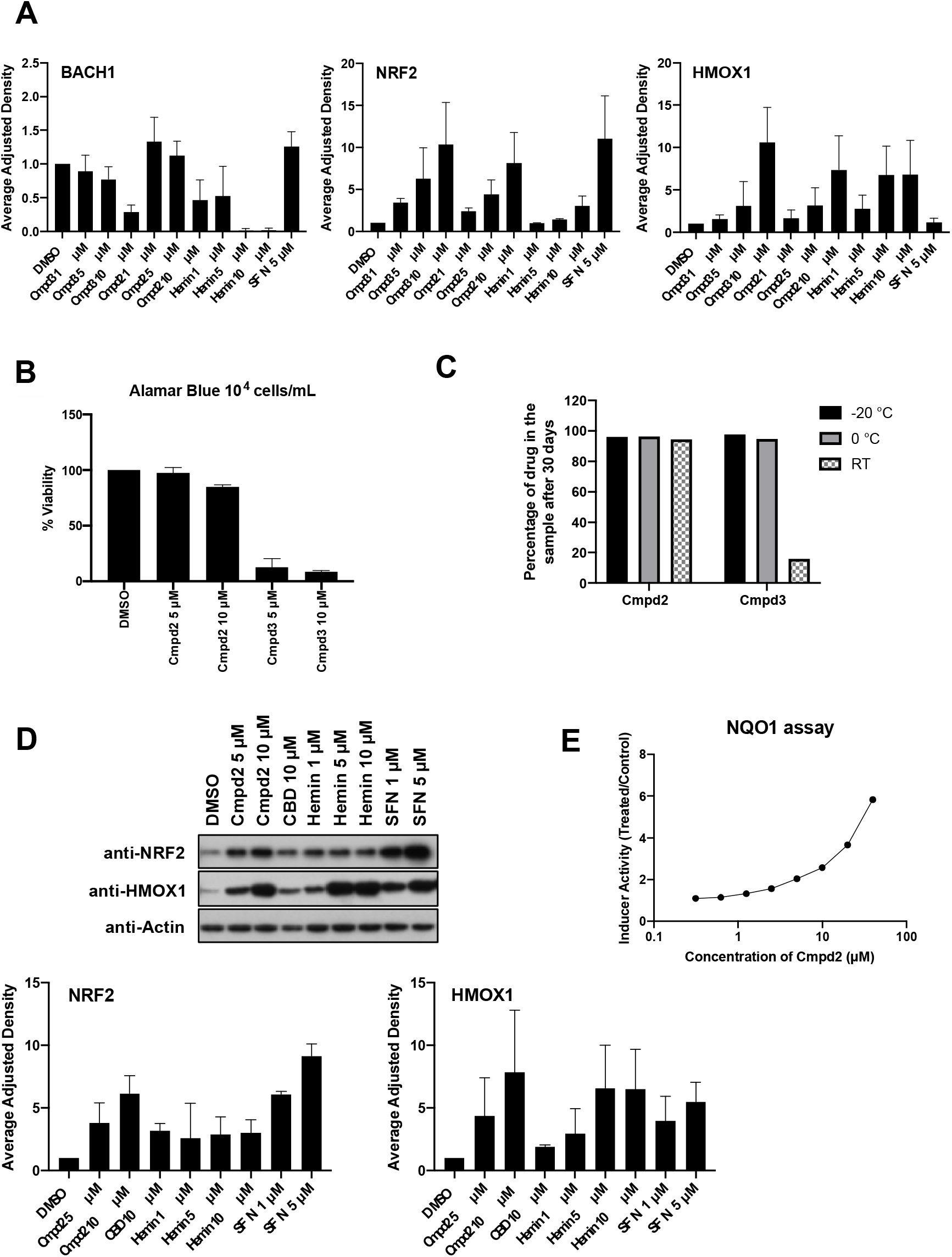
**A)** BACH1, NRF2 and *HMOX1* protein quantification against the loading control for the representative blot in Fig. 1D. Data represent means ± SD (n = 3) and are expressed relative to the DMSO treated samples. **B)** HaCaT cells were seeded in 96-well plates, and treated with DMSO or increasing concentrations of compound 3 or compound 2. 16 hours later, Alamar Blue was added to the wells and the cells were incubated for 4 hours at 37 °C. The fluorescence was measured using a plate reader and viability was calculated relative to the untreated control (n=3). *P ≤ 0.05, **P≤ 0.01, ***P ≤ 0.001 **C)** Drug samples were stored under not controlled conditions at room temperature (20-30 °C), 0 °C or −20 °C. The samples were retested after 30 days by HPLC. Percentages are expressed relative to a reference sample. **D)** Hepa-1c1c7 cells were exposed to either DMSO or increasing concentrations of compound 2, hemin or SFN for 3 hours. Cells were lysed and NRF2, *HMOX1* and Actin protein levels were analyzed by Western Blot. Upper panel is a representative western blot and bottom panels show the quantification of NRF2 and *HMOX1* protein levels against the loading control. Data represent means ± SD (n = 3) and are expressed relative to the DMSO treated samples. **E)** Hepa1c1c7 cells were treated with eight different concentrations of compound 2 in eight replicates. After 48 hours cells were lysed and the specific activity of NQO1 was analysed. The average of two independent experiments is shown.

**Suppl. Figure S2.**
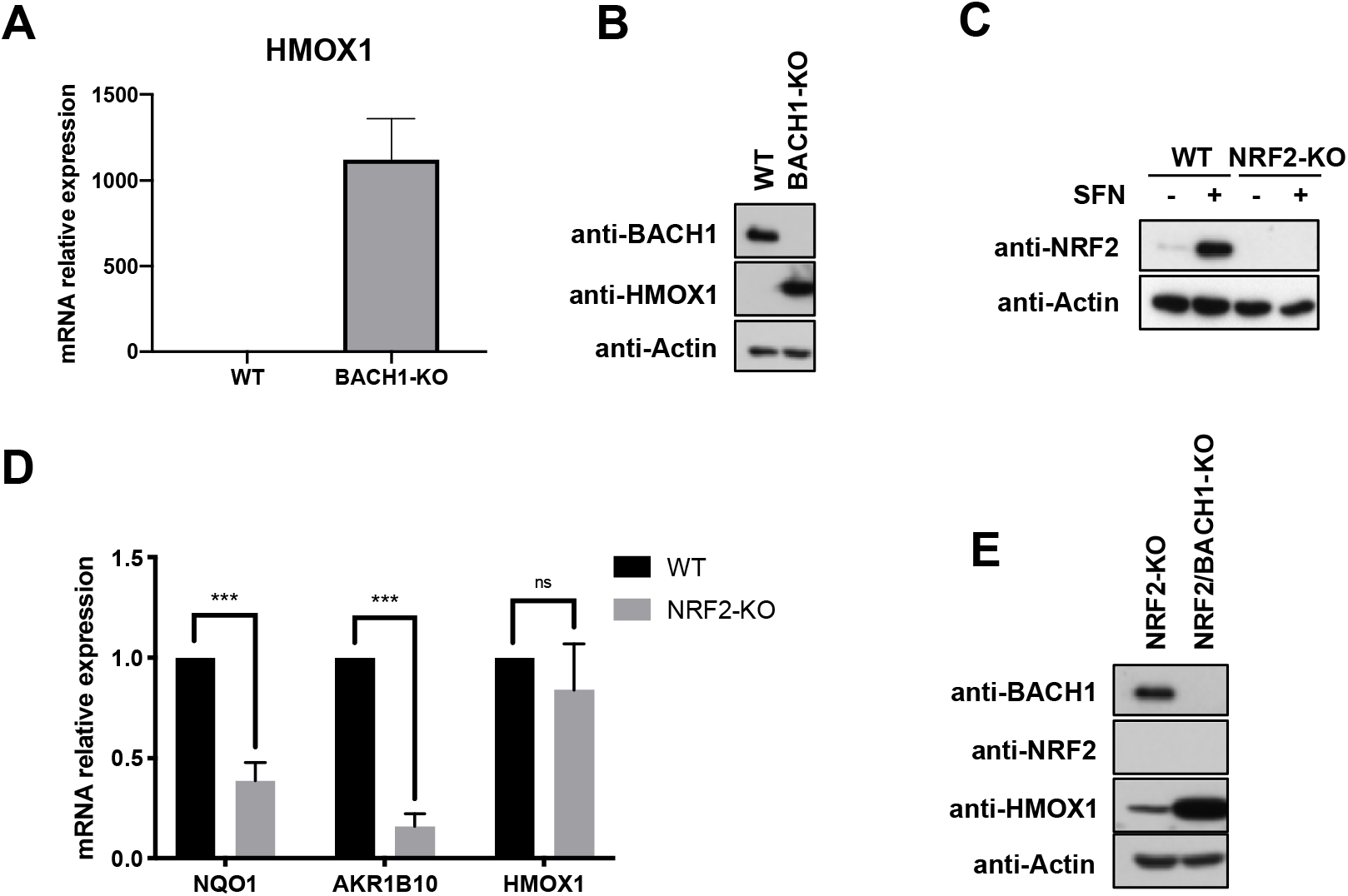
**A)** HaCaT WT and BACH1-KO cells were harvested and mRNA levels of *HMOX1* were quantified by real-time PCR. The data were normalized using HPRT1 as an internal control. Data represent means ± SD (n=3) and are expressed relative to WT sample. **P≤ 0.01 **B)** HaCaT WT and BACH1-KO cells were harvest and lysed. BACH1, *HMOX1* and Actin protein levels were analysed by Western Blot. **C)** HaCaT WT and NRF2-KO cells were incubated with either DMSO or SFN (5 μM) for 3 hours. Cells were lysated and NRF2 and Actin protein levels were analysed as previously described. **D)** HaCaT WT and NRF2-KO cells were collected and lysed. mRNA levels for NQO1, *AKR1B10* and *HMOX1* were quantified using real-time PCR (n=3). ns P ≥ 0.05, ***P ≤ 0.001. **E)** HaCaT NRF2-KO cells and HaCaT NRF2-KO/BACH1-KO cells were harvested and lysed. BACH1, NRF2, *HMOX1* and Actin protein levels were analysed by Western Blot.

**Suppl. Figure S3.**
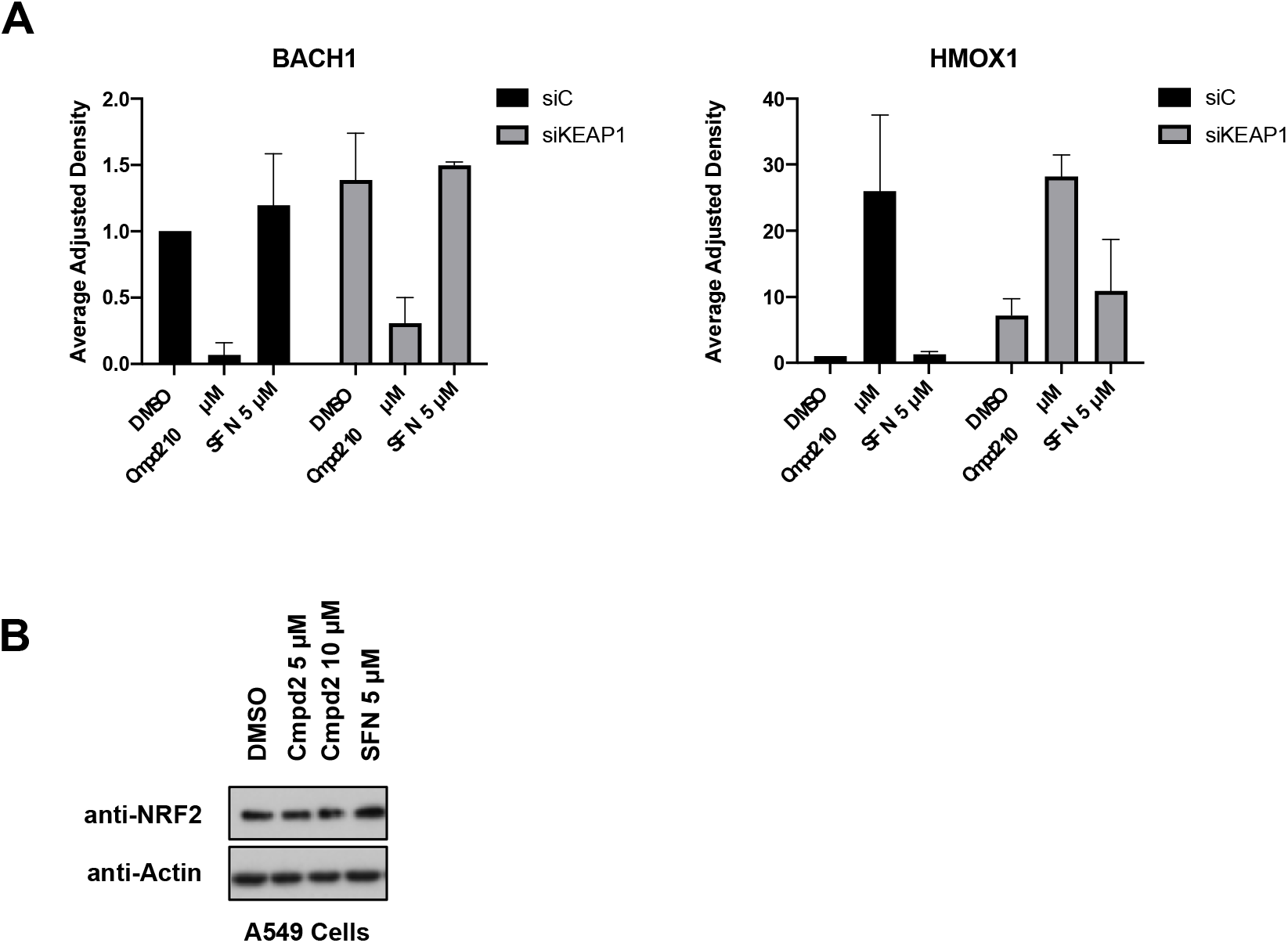
**A)** Quantification of BACH1 and *HMOX1* protein levels against the loading control from Fig. 2E. Data represent means ± SD (n = 2) and are expressed relative to the siControl DMSO sample. **B)** A549 cells were incubated with DMSO, SFN (5 μM) or increasing concentrations of compound 2 for 3 hours. Cells were lysed and NRF2 and Actin protein levels were analysed by Western Blot.

